# Evidence for low affinity of GABA at the vesicular monoamine transporter VMAT2 – implications for transmitter co-release from dopamine neurons

**DOI:** 10.1101/2024.07.04.602053

**Authors:** Fabian Limani, Sivakumar Srinivasan, Michaela Hanzlova, Ségolène La Batide-Alanore, Sigrid Klotz, Thomas S. Hnaskoxs, Thomas Steinkellner

## Abstract

**Background and Purpose:** Midbrain dopamine (DA) neurons comprise a heterogeneous population of cells. For instance, some DA neurons express the vesicular glutamate transporter VGLUT2 allowing these cells to co-release DA and glutamate. Additionally, GABA may be co-released from DA neurons. However, most cells do not express the canonical machinery to synthesize GABA or the vesicular GABA transporter VGAT. Instead, GABA seems to be taken up into DA neurons by a plasmalemmal GABA transporter (GAT1) and stored in synaptic vesicles via the vesicular monoamine transporter VMAT2. Yet, it remains unclear whether GABA indeed interacts with VMAT2, or whether another transmitter could be responsible for the observed inhibitory effects attributed to GABA.

**Experimental Approach:** We used radiotracer flux measurements in VMAT2 expressing HEK-293 cells and synaptic vesicles from rodents to determine whether GABA qualifies as substrate at VMAT2. mRNA *in situ* hybridization was employed to determine expression of VMAT2 and GAT1 transcripts in DA neurons of mouse and in human midbrains.

**Key Results:** We found that GABA reduced uptake of VMAT2 substrates in rodent synaptic vesicle preparations from striatum and cerebellum at millimolar concentrations but had no effect in VMAT2-expressing cells indicating that key components are missing in a non-neuronal system. Roughly 60 % of murine and human DA neurons in the substantia nigra express VMAT2 and GAT1 suggesting that many may be capable of co-releasing DA and GABA.

**Conclusion and Implication:** Our experiments suggest that GABA is a low-affinity substrate at VMAT2 with potential implications for basal ganglia physiology and disease.

**Bullet point summary:** *What is already known:*

- Subpopulations of dopamine neurons co-release glutamate and/or GABA.
- While glutamate is loaded into vesicles by VGLUT2, GABA co-release depends on GAT1 and VMAT2.

*What this study adds:*

- The relative affinity of GABA at VMAT2 was found to be in the millimolar range.
- Human midbrain dopamine neurons express GAT1.

*Clinical significance:*

- GABA co-release from midbrain dopamine neurons may also occur in humans.
- GABA co-release from dopamine neurons may play a role in neuropsychiatric diseases

## Introduction

The majority of dopamine (DA) neurons in the central nervous system are located in the midbrain which is grossly subdivided into substantia nigra *pars compacta* (SNc) and ventral tegmental area (VTA) (Dahlstroem et al., 1964). Though the overall number of DA neurons in these regions is comparatively small (about 500,000 DA neurons in the human SNc and VTA and about 25,000 DA neurons in mouse SNc and VTA) (Damier et al., 1999; German et al., 1996; Hirsch et al., 1989; Nelson et al., 1996), the DA system is of particular interest to neuropharmacology because of many neuropsychiatric diseases associated with DA system dysfunction, most notably Parkinson’s disease (PD), schizophrenia, substance use disorders and attention-deficit hyperactivity disorder (Carlsson, 2001; Costa et al., 2022; Hornykiewicz, 2006; Iversen et al., 2007).

For a long time, DA neurons were thought to be a rather homogenous population of cells that despite innervating different brain structures and involvement with many neurologic and psychiatric diseases, are largely similar due to their synthesis and release of the neurotransmitter DA. The complex nature and heterogeneity of the DA system was only slowly recognized in the last 30 years and became widely appreciated with advances in single-cell profiling (for summary see (Poulin et al., 2020)).

One particular feature that underlines the complexity of midbrain DA neurons results from co-release of multiple neurotransmitters. For instance, subsets of DA neurons located in the medial VTA and dorsolateral SNc express the vesicular glutamate transporter VGLUT2 and co-release glutamate, the major excitatory neurotransmitter in the central nervous system (Dal Bo et al., 2008; Kawano et al., 2006). Expression of VGLUT2 is both necessary and sufficient to allow DA neurons to store and release glutamate (Hnasko et al., 2010; Steinkellner et al., 2018; Yamaguchi et al., 2011), because glutamate is presumably present in all neurons at millimolar concentrations (Featherstone, 2010).

It was also shown that some DA neurons co-release the major inhibitory neurotransmitter ɣ-aminobutyric acid (GABA) (Tritsch et al., 2012) but GABA co-release is anatomically and functionally less well defined because DA neurons do not widely express the classical enzymes required for GABA synthesis (glutamic acid decarboxylase 1 and 2; GAD1/2) and/or the vesicular GABA transporter (VGAT), which is generally required for vesicular sequestration of GABA into synaptic vesicles (McIntire et al., 1997).

Rather, the vesicular transporter for DA, VMAT2, seems to be required for the transport of GABA into synaptic vesicles (Tritsch et al., 2012). This finding was rather surprising because GABA is a zwitterionic amino acid and structurally very different from classical VMAT2 substrates that have an aromatic ring and a positive charge (Peter et al., 1994; Yelin et al., 1995; Zheng et al., 2006). In contrast to glutamate, which is presumably present in all cells with active gene expression, GABA is typically produced through decarboxylation of glutamate in cells containing GAD1 or GAD2, which are little expressed in DA neurons (Azcorra et al., 2023; Gaertner et al., 2024; Tritsch et al., 2014). But how do DA neurons get GABA in the first place? It seems that GABA is largely taken up into DA neurons from outside via the plasmalemmal GABA transporters GAT1 (Tritsch et al., 2016). Alternatively, GABA may be synthesized *de novo* in DA neurons through aldehyde dehydrogenase 1a1 (ALDH1a1) (Kim et al., 2015), an enzyme widely expressed in SNc DA neurons.

Together, though there is strong electrophysiological and genetic evidence that GABA is released from DA neurons in a VMAT2-dependent manner, the relative affinity of GABA at VMAT2 remains unknown. Using radiotracer flux measurements, we show that GABA weakly competes with monoamine uptake at VMAT2 that is considerably lower than that of the cognate substrates DA or serotonin (5-HT). Finally, we provide evidence for widespread expression of GAT1 not only in mouse but also in human midbrain DA neurons suggesting that mechanisms subserving GABA co-release may be conserved across mice and humans.

## Materials and Methods

### Materials

3,4-[Ring-2,5,6-^3^H]-dihydroxyphenylethylamine (dopamine), 1 mCi/mL and 5-[1,2-^3^H(N)]-hydroxytryptamine creatinine sulfate (serotonin; 5-HT), 1 mCi/mL were purchased from Revvity. (±)-alpha-[2-^3^H]-dihydrotetrabenazine, 10-20 Ci/mmol was purchased from American Radiolabeled Chemicals. Scintillation fluid (Rotiszint eco plus) was purchased from Carl Roth GmbH (Karlsruhe, Germany). Tetrabenazine was purchased from Eubio. Rabbit GFP polyclonal antiserum (A-11122; RRID AB_221569) is from Thermo Fisher Scientific. RNAscope Multiplex Fluorescent Reagent Kit v2, RNAscope 2.5 HD Duplex Reagent Kit, mouse RNAscope probes against VMAT2 (Mm-Slc18a2-C3; # 425331-C3), GAT1 (Mm-Slc6a1-C2; # 444071-C2) and VGLUT2 (Mm-Slc17a6; # 319171-C1), and human probes (Hs-TH-C2, #441651-C2; Hs-Slc6A1, # 545121; Hs-Slc17A6, #415671) were purchased from ACDBio/Bio-Techne Ireland Limited. All other chemicals were purchased from Sigma-Aldrich (Vienna, Austria).

### Animals

We used adult male and female C57BL/6J mice and adult female Sprague-Dawley rats (female rats were used because they were specifically ordered from our Animal Breeding Facility as pregnant dams to harvest pups for primary neuronal cell culture). Mice and rats were group-housed, and maintained on a 12:12-hour light: dark cycle with food and water available *ad libitum*. All animals were used in accordance with protocols approved by the Animal Welfare Committee of the Medical University of Vienna and the Austrian Federal Ministry of Science and Research (BMBWF licenses 2021-0.373.073 and 2023-0.515.074).

### Human tissue

De-identified formalin-fixed paraffin-embedded sections (FFPE) from the midbrain including segments of the substantia nigra at the level of the emergence of the third cranial nerve of two adult patients (one male, 61 years and one female, 76 years) without pathological changes in the substantia nigra were obtained from the neuro biobank of the Division of Neuropathology and Neurochemistry/Department of Neurology at the Medical University of Vienna.

### Cell culture

HEK-293 cells were purchased from ATCC (CRL-1573, ATCC, Manassas, VA, USA; RRID: CVCL_0045) and transfected with human VMAT2-eGFP in pcDNA6.2 using jetPRIME (Polyplus). A polyclonal cell line was generated through blasticidin S (Thermo Fisher Scientific; 10 µg/mL) selection and maintained in Dulbecco’s modified eagle medium (DMEM; Sigma-Aldrich) containing 10 % fetal bovine serum (Sigma-Aldrich), penicillin (100 IU/mL, Sigma-Aldrich), streptomycin (100 µg/mL, Sigma-Aldrich) and blasticidin S (6 µg/mL). Cells were maintained in a humidified 5 % CO_2_ atmosphere at 37°C.

### Fluorescent microscopy

Live VMAT2-eGFP polyclonal cells were imaged at a Nikon confocal microscope using a 100x objective. Cells were incubated with 0.05 % trypan blue in phosphate-buffered saline (PBS) for 10 min prior to imaging.

### Immunoblotting

VMAT2-eGFP Cells were grown in a 10 cm plastic dish, washed two times in PBS and lysed in RIPA buffer containing (50mM Tris.HCl, 150mM NaCl, 1mM EDTA, 1% Triton X-100, 0.1% SDS and 1% deoxycholate supplemented with protease inhibitors) and incubated at 4°C for 30 min on a rotator followed by centrifugation at 12,600 x g for 10 min.

Proteins were separated on a 10 % SDS-PAGE and electrotransferred onto nitrocellulose before incubation with polyclonal GFP antiserum (A-11122; Thermo Fisher Scientific) overnight. IRDye 680-RD-labeled secondary antibody was obtained from LI-COR Biotechnology GmbH (Bad Homburg, Germany). and visualized using the LI-COR Odyssey CLx infrared imaging system.

### Synaptic vesicle preparation

Vesicles were prepared as described previously (Pifl et al., 2014) with minor modifications. Briefly, mice were deeply anaesthetized with pentobarbital (100 mg/kg i.p.; Exagon^®^ 500 mg/mL, Richter Pharma) before decapitation. Rats were euthanized by CO_2_ asphyxiation and then decapitated. Mouse and rat brains were rapidly removed and striatum and cerebellum were dissected. Rodent striatum and cerebellum were homogenized in ice-cold 0.3 M sucrose containing 25 mM Tris (pH = 7.4) using a glass Teflon Potter-type homogenizer (10-15 strokes). Homogenates were centrifuged for 15 minutes at 1000 x g at 4°C. The supernatant was centrifuged at 20,000 x g for 30 min at 4°C. The resulting ‘P2’ pellet was osmotically shocked through resuspension in 2 mL ice-cold H_2_O and additional 5-10 strokes in a glass Teflon Potter-type homogenizer, the supernatant (‘SN2’) was kept on ice. The aqueous P2 suspension was centrifuged at 22,000 x g for 15 min and osmolarity of SN re-adjusted by addition of 1.3 M potassium phosphate buffer (pH=7.4) in 1/10 of the volume. SN2 (from above) was centrifuged at 100,000 x g for 30 min at 4°C and resuspended in the hyposomotically shocked P2 suspension. The combined suspension was centrifuged at 100,000 x g for 60 min at 4°C and the resulting “LP2” pellet was resuspended in 0.13 M potassium phosphate (KP) buffer and kept on ice or stored at -80°C until the start of the experiment.

### Radioligand uptake

Polyclonal HEK-293-VMAT2:eGFP cells were grown in a 48-well plate coated with poly-D-lysine (100 µg/mL). Prior to uptake, cells were washed with 200 µL NMDG buffer (in mM: 150 N-methyl-D-glucamine, 10 HEPES, 2 magnesium sulfate, 2 potassium chloride, 10 potassium gluconate; pH = 7.3) and permeabilized in 50 µM digitonin in NMDG buffer for 15 minutes at room temperature. Preincubation with inhibitors (reserpine, tetrabenazine) or substrates (dopamine) was performed for 5 minutes in NMDG buffer followed by incubation containing 100 nM [^3^H]dopamine (40 Ci/mmol; Perkin-Elmer) in addition to unlabeled substrates or inhibitors.

Uptake was terminated after 30 minutes at room temperature by addition of ice-cold NMDG buffer. After two additional washes with ice-cold NMDG buffer, cells were lysed in 1 % SDS, transferred to scintillation vials containing scintillation cocktail and radioactivity was measured in a liquid scintillation counter. Each cell uptake was performed in triplicate determinations.

Uptake in rodent or human synaptic vesicles was performed in KP buffer containing 2 mM ATP-Mg and 2 mM potassium chloride in a final volume of 200 µL and in the presence of [^3^H]5-HT (100 nM). Uptake was started after addition of 10 µL vesicles (roughly 2 µg of protein) to ^3^H solution and incubation in a water bath at 30°C for 5 minutes. Ice-cold KP buffer was added to tubes to terminate the uptake followed by rapid filtration of suspension through polyethyleneimine-coated (1 % w/v) GF/B glass fiber filters (Whatman) and two additional washes with 2 mL KP buffer. Filters were then transferred to scintillation vials, vortexed and measured in a liquid scintillation counter (Perkin Elmer/Revvity). Each vesicle uptake was performed in duplicate determinations.

### Radioligand binding

Binding of [^3^H]-dihydrotetrabenazine was performed as described (Pifl et al., 2014). Briefly, striatal synaptic vesicles were added to binding buffer (25 mM sodium phosphate; pH = 7.4) containing 20 nM [^3^H]-dihydrotetrabenazine and different concentrations of unlabeled GABA. The reaction was incubated for 90 min at 30°C. Non-specific binding was determined in the presence of 10 µM tetrabenazine. Binding was stopped by adding ice-cold binding buffer and samples were filtered onto GF/B filters presoaked in 1 % polyethylenimine using an automated cell harvester filtration device (Skatron Instruments AS). The radioactivity bound to filters was measured by liquid scintillation counting. Binding reactions were always determined in duplicates.

### Fluorescent mRNA in situ hybridization in mouse brain

Mice were anesthetized with pentobarbital and killed by decapitation. Brains were extracted rapidly, frozen in chilled isopentane and stored at -80°C. Sections were serially cut (20 μm) on a cryostat and mounted directly onto Epredia Superfrost Plus glass slides (Epredia Netherlands B.V.). Slides were stored at -80°C until starting the multiplex fluorescent RNAscope assay v2 (Advanced Cell Diagnostics). Briefly, sections were fixed with 4% PFA for 1 hour at 4°C followed by dehydration in increasing ethanol concentrations and protease IV treatment. RNA hybridization probes included antisense probes against VMAT2 (Mm-Slc18a2-C3; # 425331-C3), GAT1 (Mm-Slc6a1-C2; # 444071-C2) and VGLUT2 (Mm-Slc17a6; # 319171-C1).

Slides were counterstained with DAPI and coverslipped using Fluoromount-G mounting medium. Images were taken at 20 x magnification using a Vectra Polaris slide-scanner (Akoya) at the imaging core facility of the Medical University of Vienna.

### Chromogenic mRNA in situ hybridization in human brain

Slides containing human FFPE midbrain sections were baked at 60°C for 1 h in HybEZ II oven (ACDBio) and deparaffinized in xylenes followed by washes in 100% ethanol and air dried for 15 min. Next, tissue was pretreated with H_2_O_2_, boiled in target retrieval solution and incubated with protease plus before probe hybridization (Hs-TH-C2, #441651-C2; Hs-Slc6A1, # 545121; Hs-Slc17A6, #415671) and amplification according to the RNAscope 2.5 HD Duplex Assay (ACDBio). Slides were counterstained with Mayer’s hematoxylin and coverslipped using VectaMount media. Images were taken at 40 x magnification using a Vectra Polaris slide-scanner (Akoya) at the imaging core facility of the Medical University of Vienna.

### Data analysis and statistics

Vesicle uptake and binding experiments were analyzed as follows: amount of radioactivity measured (counts per minute [cpm]) in the absence of vesicles (‘filter blank’; unspecific binding of ^3^H to the filter) was subtracted from cpm measured in the presence of vesicles containing vehicle (control) or test compounds (*i.e.* GABA, reserpine, glycine, taurine, tetrabenazine). After subtraction, average of the uptake/binding cpm obtained per experiment (each experiment was performed in duplicates) in the presence of vehicle (control) was set to 100 %. cpm in the presence of test compounds (GABA, reserpine, glycine, taurine, tetrabenazine) are expressed as percentage of control.

Uptakes in HEK-293 cells were analyzed the same way except that there was no ‘filter blank’ subtracted.

Cell counting: For RNAscope, a neuron was deemed positive for a given mRNA if at least 4 puncta were present in close proximity to a DAPI- (mice) or hematoxylin (humans) labeled nucleus. 4 sections covering the mouse midbrain (spaced approximately 100-150 μm) from 3 animals were counted. For human RNAscope, one section containing both hemispheres from one male and one female donor was counted. Cell fractions were averaged across the different mice (n=3) or humans (n=2) and expressed as percentage of total cell counts.

GraphPad Prism (GraphPad Software Inc., San Diego, CA) was used to analyze data and create graphs. All data are expressed as means ± SEM.

## Results

### GABA does not inhibit [^3^H]dopamine uptake in HEK-293 cells stably expressing human VMAT2

To test whether GABA would interact with VMAT2 we created a stable polyclonal HEK-293 cell line expressing human VMAT2 with a C-terminal eGFP tag (HEK-293-VMAT2:eGFP). Live cell imaging shows that VMAT2 is diffusely expressed in intracellular compartments consistent with previous reports (Erickson et al., 1992) (**Figure 1A**) Immunoblotting confirms expression of VMAT2:eGFP at around 80-90 kD (**Figure 1B**).

**Figure 1:**
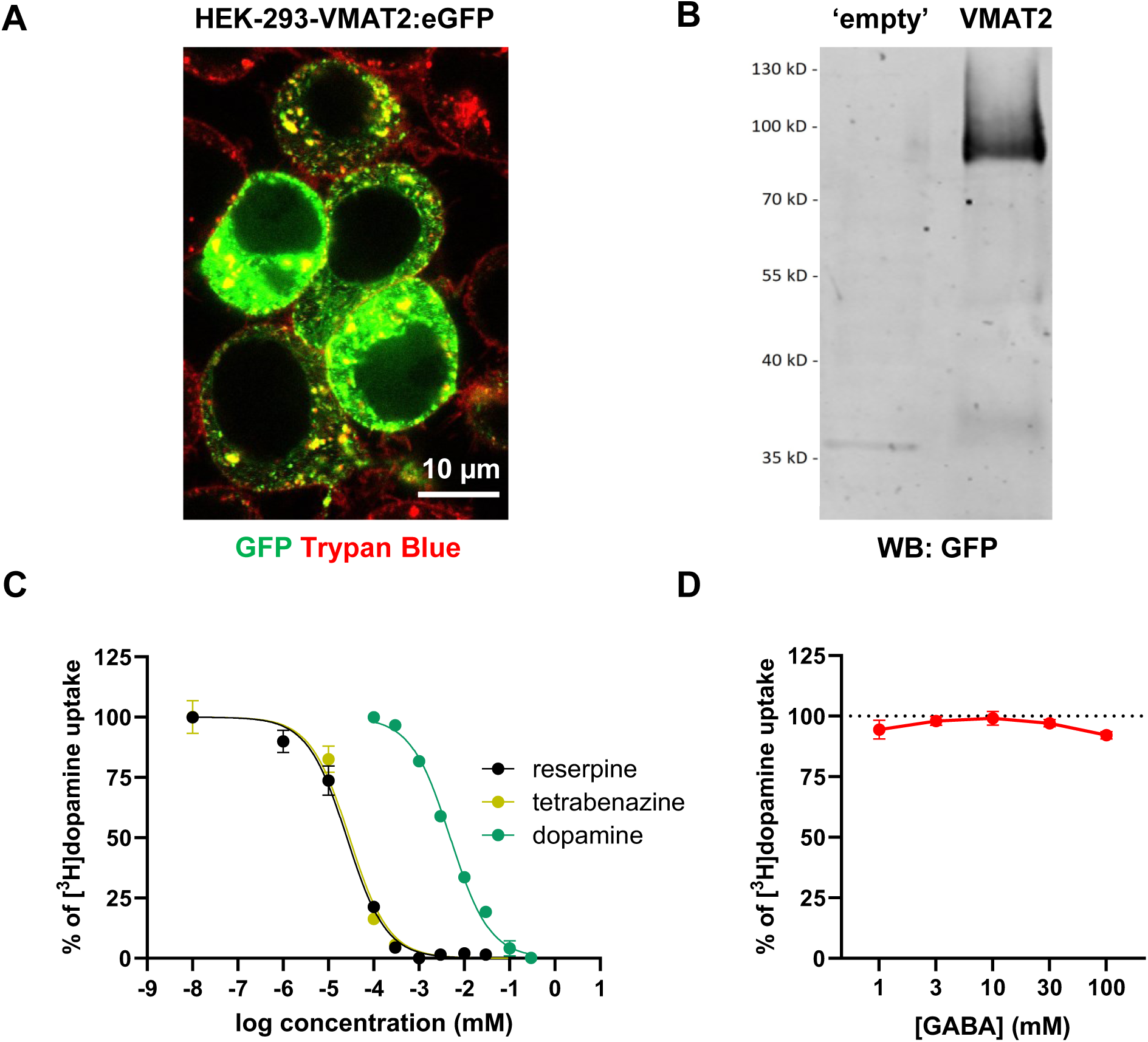
GABA did not affect dopamine uptake at VMAT2 in polyclonal HEK-293 cells. **(A)** Trypan blue labeling indicates that VMAT2:eGFP is expressed in intracellular compartments of a polyclonal HEK-293 cell line. **(B)** Detection of VMAT2:eGFP at 80-90 kD by immunoblotting using an anti-GFP antiserum. **(C, D)** Inhibition uptake of [^3^H]dopamine in digitonin-permeabilized HEK-293 VMAT2:eGFP cells with increasing concentrations of reserpine (n=2), tetrabenazine (n=2) and dopamine (n=2) **(C)** and GABA, n=8 **(D)**.

We generated this line to perform vesicular uptake assay in adherent cells. To do so we utilized the steroidal saponin detergent digitonin to permeabilize the plasma membrane in order for VMAT2 substrates to reach the intracellular compartments. Our uptake inhibition experiments demonstrate that uptake of the VMAT2-substrate [^3^H]dopamine (DA) into digitonin-permeabilized HEK-293-VMAT2:eGFP cells is potently inhibited by the competitive VMAT inhibitor reserpine (IC_50_ = 25.2 nM; 95% CI: 20.4 to 31.2 nM; n=2), the non-competitive inhibitor tetrabenazine (IC_50_ = 28.8 nM; 95% CI: 21.4 to 38.7 nM; n=2) as well as DA itself (IC_50_ = 4.9 µM; 95% CI: 4.4 to 5.6 µM; n=2) (**Figure 1C**). These affinities are well in line with published values (Erickson et al., 1992; Kanner et al., 1979; Liu et al., 1992) and support the robustness of this assay to investigate VMAT2 function in a heterologous expression system.

Accordingly, we used these cells to test whether GABA would reduce uptake of [^3^H]DA into HEK-293-VMAT2:eGFP cells as would be expected if it were a VMAT2 substrate. However, GABA did not concentration-dependently affect [^3^H]DA uptake (**Figure 1D**).

### GABA weakly inhibits uptake of [^3^H]5-HT in rodent striatal synaptic vesicles

HEK-293 cells are non-neuronal cells and do not have synaptic vesicles. Expression of VMAT2:eGFP in intracellular compartments of HEK-293 cells may therefore not well mimic the native environment of VMAT2 present in neuronal cells such as DA neurons. We therefore tested effects of GABA at VMAT2 in isolated synaptic vesicles from rodent brains. To measure VMAT2 uptake in vesicles we relied on [^3^H]serotonin (= 5-hydroxytryptamine; 5-HT) as a substrate rather than [^3^H]DA because of its superior signal-to-noise ratio, which may be related to 5-HT’s slightly higher affinity at VMAT2 (Erickson et al., 1992; Liu et al., 1992). In our preparation, 5-HT shows saturable uptake kinetics with an apparent Michaelis-Menten constant (K_M_) of about 200 nM (**Figure 2A**).

**Figure 2:**
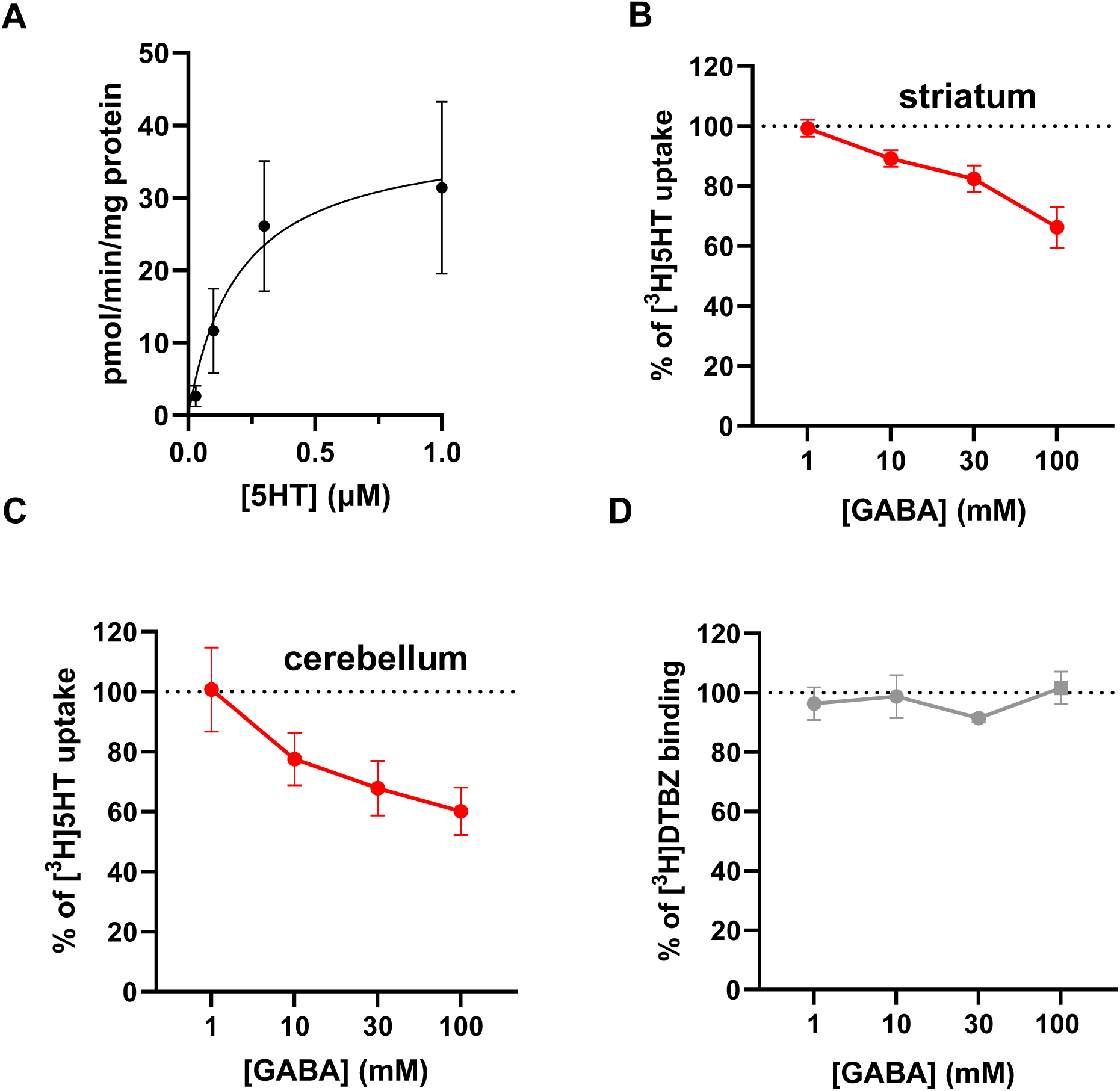
GABA decreased uptake of 5-HT in mouse synaptic vesicles. **(A)** Saturation uptake of [^3^H]5-HT in mouse striatal synaptic vesicles (n=5). **(B, C)** GABA concentration-dependently decreases [^3^H]5-HT uptake in striatal (n=5-9) **(B)** and cerebellar (n=5) **(C)** vesicles. **(D)** Binding of [^3^H]dihydrotetrabenazine is not affected by GABA in striatal synaptic vesicles (n=5).

In contrast to what we observed in HEK293-VMAT2:eGFP cells, GABA reduced uptake of [^3^H]5-HT into both, mouse and rat striatal synaptic vesicles at millimolar concentrations (**Figure 2B** and **Suppl. Figure 2A**). This suggests that GABA may indeed be a low-affinity substrate at VMAT2.

The rodent striatum receives very dense axonal input from midbrain DA neurons, and comparatively fewer input from serotonergic or noradrenergic fibers suggesting that the preponderance of VMAT2 measured in striatal synaptic vesicles is derived from DA neurons, although a separation between the different vesicles is of course impossible. Based on neurochemical measurements of tissue monoamine levels by high-performance liquid chromatography, there is about 10-fold more DA than 5-HT, and about 10-fold more 5-HT than norepinephrine (NE) (Peneder et al., 2011).

Nonetheless, we wondered whether VMAT2 would also be GABA-sensitive in a brain area where VMAT2 derives primarily from serotonergic and noradrenergic innervation and less from DA neurons. Hence, we also prepared synaptic vesicles from mouse cerebellum, which has 10-40 fold lower levels of DA compared to NE (Glaser et al., 2006) and again measured [^3^H]5-HT uptake in the presence of increasing concentrations of GABA. Comparable to our findings in mouse and rat striatal vesicles, cerebellar [^3^H]5-HT uptake was decreased in the presence of millimolar concentrations of GABA (**Figure 2C**).

Next, we wondered whether GABA would displace the high-affinity non-competitive antagonist [^3^H]dihydrotetrabenazine (DTBZ) from binding to VMAT2. The apparent dissociation constant (K_D_) of [^3^H]DTBZ in mouse striatal synaptic vesicles was determined at around 5 nM (**Supplemental Figure 1**). Striatal synaptic vesicles were incubated with a saturating concentration of [^3^H]DTBZ (20 nM) and increasing concentrations of unlabeled GABA. However, we did not observe any [^3^H]DTBZ displacement even at very high millimolar GABA concentrations (100 mM) in both mouse (**Figure 2D**) and rat striatal vesicles (**Supplemental Fig. 2B**).

### Effects of glycine and taurine on VMAT2

It is possible that not GABA but another inhibitory transmitter is packaged into vesicles by VMAT2 and responsible for the activation of postsynaptic striatal GABA_A_ receptors that was reported after optogenetic stimulation of DA terminals (Melani et al., 2022; Patel et al., 2024; Tritsch et al., 2012; Tritsch et al., 2014). We tested two such potential compounds in the mouse vesicle uptake, glycine and taurine, both of which are structural analogues of GABA that have been reported to activate GABA_A_ receptors, though presumably with much weaker affinity (Jonas et al., 1998).

While glycine did not affect uptake of [^3^H]5-HT into mouse striatal synaptic vesicles even at high millimolar concentrations (**Figure 3A**), taurine weakly reduced [^3^H]5-HT uptake into mouse striatal vesicles at 10 and 30 mM (**Figure 3B**) suggesting that it may also be packaged into vesicles by VMAT2. Importantly, the lack of any inhibitory effects of glycine even at the high concentration of 100 mM argues against simple unspecific osmotic effects inhibiting substrate uptake at VMAT2 in our assay.

**Figure 3:**
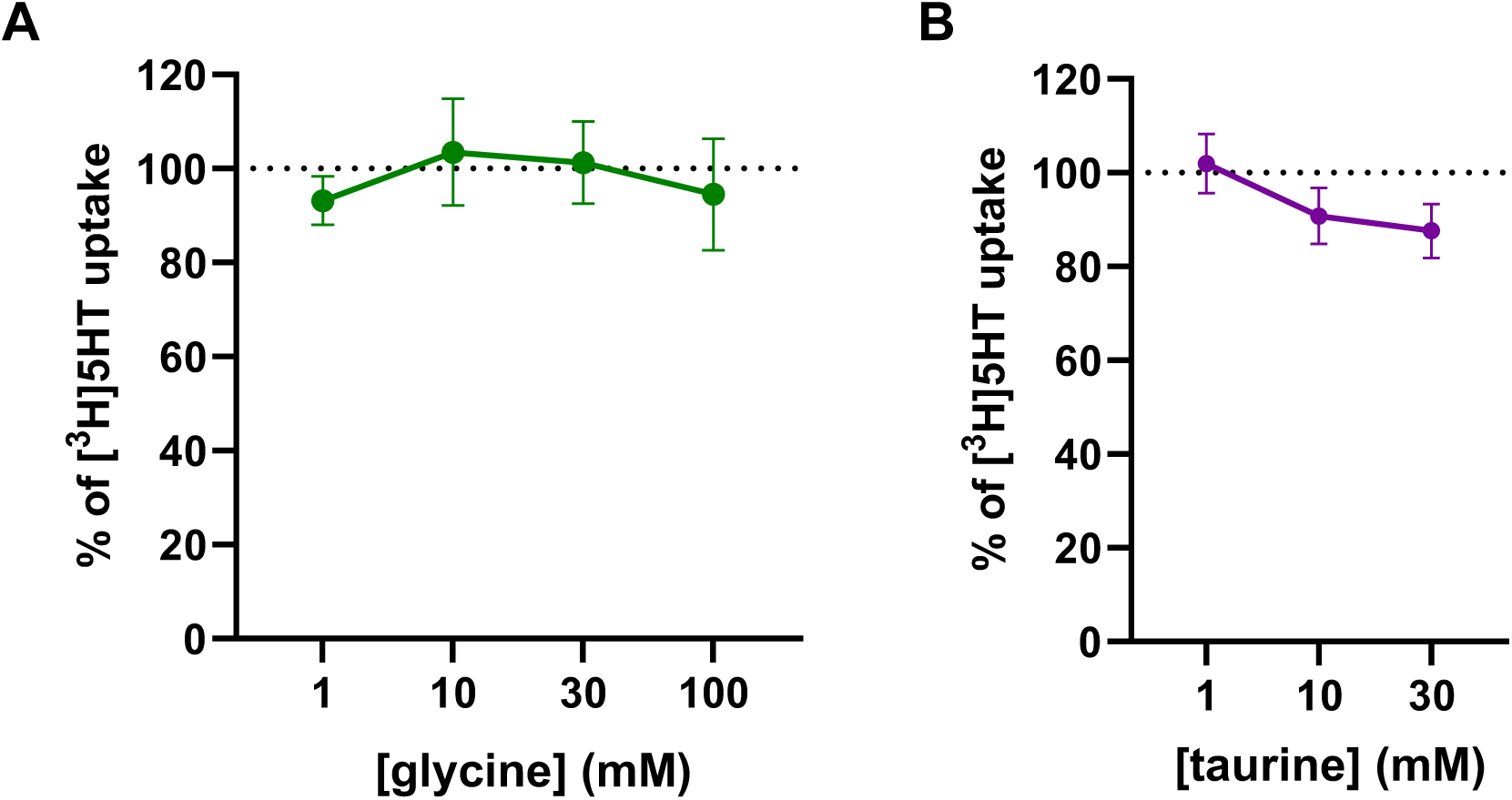
Taurine but not glycine reduces 5-HT uptake in mouse synaptic vesicles. Effects of **(A)** glycine (n=5) and **(B)** taurine (n=6) on [^3^H]5-HT uptake in mouse striatal synaptic vesicles.

### Gat1 mRNA is expressed in most SNc and VTA DA neurons and partially overlaps with Vglut2

GAT1 was proposed to be mainly responsible for GABA uptake into DA neurons and found to be expressed in almost 90 % of VMAT2^+^ DA neurons in the mouse SNc (Tritsch et al., 2014).

If VMAT2 transports GABA, all GAT1^+^ DA cells are theoretically equipped with the machinery to store and release GABA. We wondered whether GAT1 was only expressed in SNc or also in VTA DA neurons, and whether GAT1^+^ DA neurons would be distinct from the VGLUT2^+^ DA neuron population.

To investigate this, we performed multiplex fluorescent *in situ* hybridization in mouse midbrain sections targeting *Slc18a2* (*VMAT2*), *Slc3a1* (*GAT1*) and *Slc17a6* (*VGLUT2*) mRNAs (**Figure 4A-B**). We determined that about 60 % of SNc DA neurons were positive for *VMAT2* and *GAT1* mRNAs (**Figure 4C**), which is lower than previously reported (Tritsch et al., 2014). In contrast, only about 30 % of VTA DA neurons were co-expressing *VMAT2* and *GAT1* mRNAs (**Figure 4D**). In line with previous results, about 9 % of SNc and about 14 % of VTA DA neurons were positive for *VMAT2* and *VGLUT2* (**Figure 4**) (Conrad et al., 2024; Steinkellner et al., 2018).

**Figure 4:**
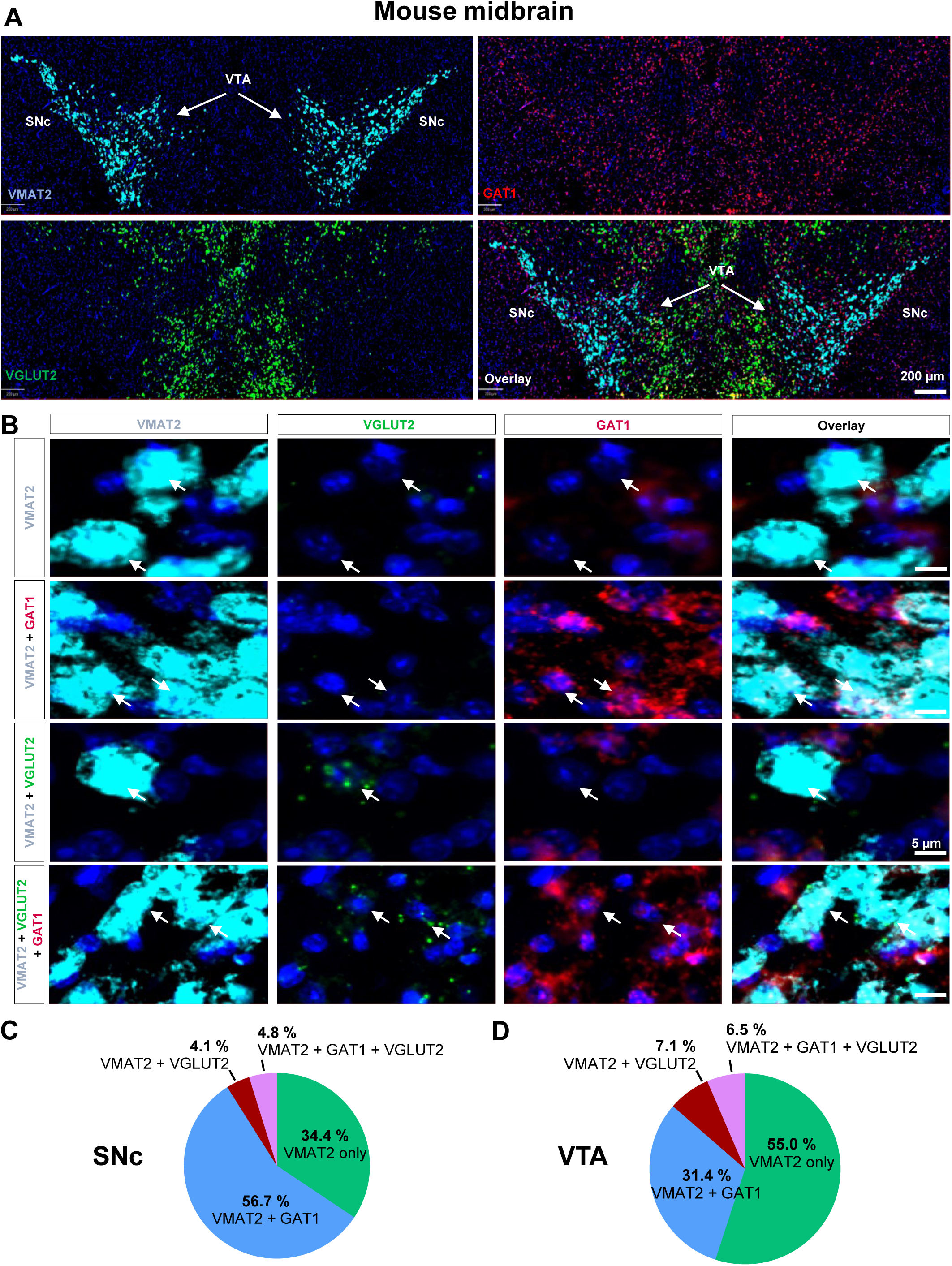
Expression of VMAT2, GAT1 and VGLUT2 mRNAs in mouse ventral midbrain dopamine neurons. **(A)** Example *in situ* hybridization images in coronal sections through the mouse ventral midbrain; mRNA encoding VMAT2 (light blue), GAT1 (red) and VGLUT2 (green) with DAPI counter-stain (dark blue). **(B)** Higher power images of different cell-types in the SNc. White arrows indicate cell-types highlighted on the left. **(C, D)** Pie charts quantifying the relative percentages of single, double or triple positive cells in SNc **(C)** and VTA **(D)**. n=3.

Interestingly, about half of these *VMAT2^+^*/*VGLUT2^+^* cells in both, SNc and VTA also expressed *GAT1* (4.8 %) (**Figure 4C**) suggesting that these neurons may co-release DA, glutamate and GABA.

### Human DA neurons express Gat1 mRNA and may co-release GABA

Human DA neurons share many molecular features with their mouse counterparts. For instance, the expression pattern of VGLUT2 in DA neurons seems similar in mice and humans (Buck et al., 2021; Root et al., 2016; Steinkellner et al., 2022) suggesting that human DA neurons are capable of glutamate co-release. But it is not known whether human midbrain DA neurons express GAT1, or whether they may co-release GABA. We used postmortem human midbrain sections from one male (61 years) and one female (76 years) donor and performed chromogenic duplex mRNA *in situ* hybridization to detect human *tyrosine hydroxylase* (*TH*), a catecholaminergic marker, and *GAT1* mRNAs (**Figure 5**). Similar to our observations in mice, about 66 % of human SNc (**Figure 5C**) and about 55 % of human VTA (**Figure 5D**) DA neurons were positive for *GAT1* mRNA. This suggests that human SNc and VTA DA neurons may co-release GABA.

**Figure 5:**
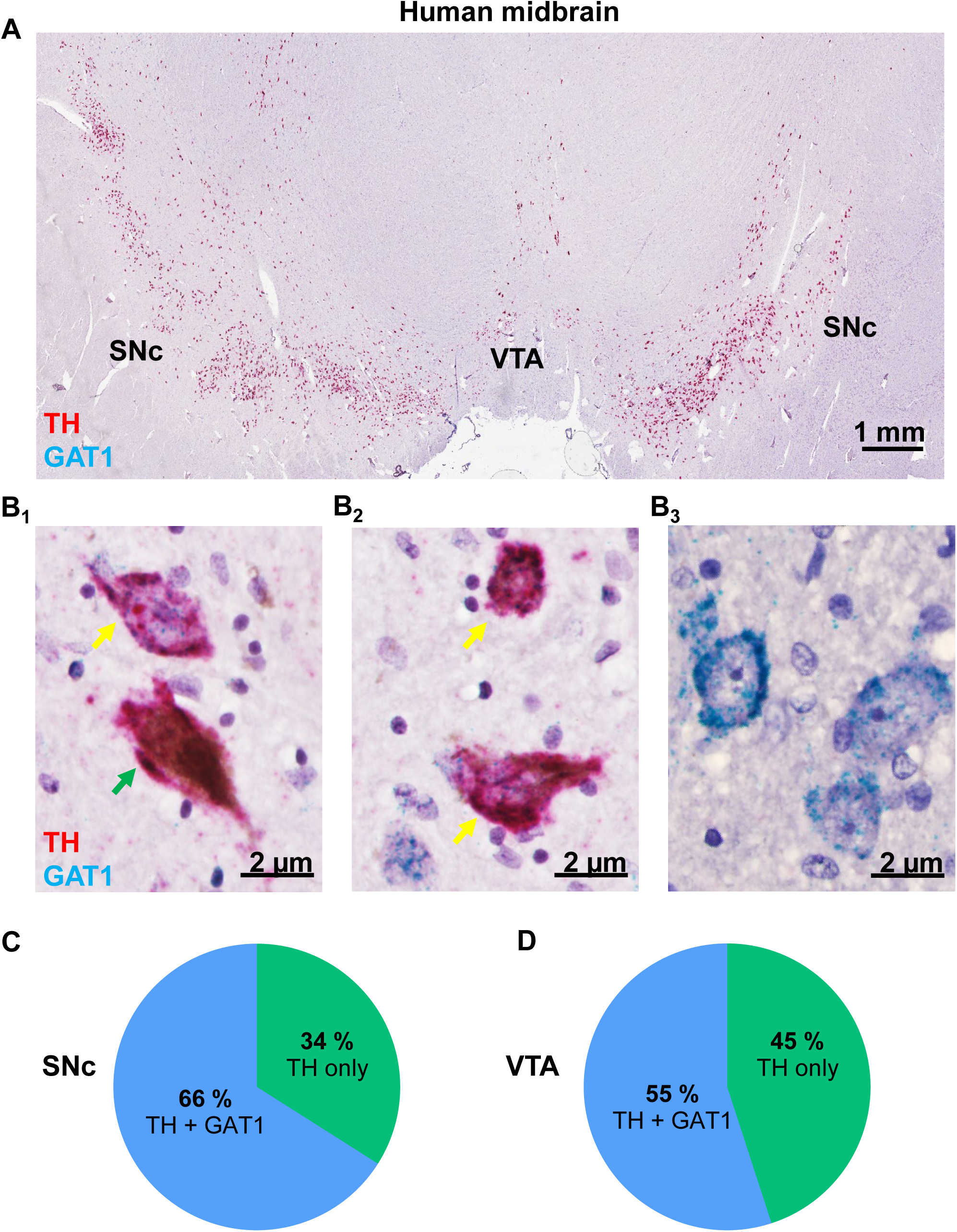
GAT1 is expressed in human SNc and VTA DA neurons. **(A)** Widefield view of transverse section containing SNc from a human brain donor with chromogenic labeling for mRNAs encoding TH (diffuse magenta) and GAT1 (blue puncta) counter-stained with hematoxylin; brown signal is neuromelanin. **(B)** Higher magnification images of SNc DA neurons (B_1_ and B_2_) expressing TH with (yellow arrow) or without (green arrow) GAT1, or putative GABA neurons expressing GABA without TH in the SNc (B_3_). **(C, D)** Pie charts quantifying the relative percentages of single, double or triple positive cells in SNc **(C)** and VTA **(D)**. n=2.

## Discussion

The heterogeneity of midbrain DA neurons is increasingly recognized and may help better explain and understand the manifold diseases associated with DA dysfunction such as Parkinson’s disease, schizophrenia, substance use disorders, or ADHD.

One interesting feature that highlights the complexity of midbrain DA neurons is their capability to co-release transmitters other than DA. The old idea that one neuron releases just one neurotransmitter (‘Dale’s principle’) turns out to be the exception rather than the rule within the DA system, but possibly also in other neural circuits (Trudeau et al., 2014; Wallace et al., 2023). As for the ventral midbrain DA system, it is now well established that the excitatory neurotransmitter glutamate is co-released from subpopulations of VTA and SNc DA neurons, because these neurons express VGLUT2, which is both necessary and sufficient to store and release glutamate (Adrover et al., 2014; Chuhma et al., 2004; Hnasko et al., 2010; Silm et al., 2019; Stuber et al., 2010). In fact, the only requirement for any neuron including DA neurons to co-release glutamate seems to be the expression of a vesicular glutamate transporter (Steinkellner et al., 2018; Takamori et al., 2000), because glutamate is estimated to be present at millimolar concentrations in all neurons (Featherstone, 2010).

In contrast, co-release of GABA from DA neurons is less well understood. Though there seem to be small subsets of DA neurons expressing GAD1/2 (Azcorra et al., 2023; Gaertner et al., 2024) and even VGAT (Conrad et al., 2024) the majority does not. Rather, it was suggested that GABA is taken up from outside by the plasmalemmal GABA transporter GAT1 (Melani et al., 2022; Tritsch et al., 2014) and/or synthesized *de novo* by ALDH1a1 (Kim et al., 2015) prior to being loaded into vesicles by VMAT2 (Melani et al., 2022; Tritsch et al., 2012).

VMAT2 is the major VMAT isoform expressed in the CNS and responsible to transport monoamines (DA, 5-HT, NE, epinephrine, histamine) into synaptic vesicles (Eiden et al., 2011). In contrast to monoamines, GABA is an amino acid that occurs as a zwitterion at physiological pH, and thus does not resemble the classical VMAT2 substrates that have an aromatic ring and a positively charged amino group (Peter et al., 1994; Yelin et al., 1995; Zheng et al., 2006).

It is therefore not surprising that GABA has never been recognized as a substrate at VMAT2. There is however now compelling pharmacological and genetic evidence that GABA can be loaded into vesicles by VMAT2. GABA co-release was reduced in the presence of VMAT2 inhibitors, and ectopic expression of VMAT2 in non-dopamine GABA neurons in which VGAT was genetically deleted was sufficient to rescue GABA release (Tritsch et al., 2012).

In the current study, we aimed to determine the relative affinity of GABA at VMAT2 and found that GABA may indeed qualify as a weak substrate at VMAT2: millimolar GABA concentrations were required to partially block uptake of the endogenous VMAT2 substrate 5-HT in rodent synaptic vesicle preparations. But does such a low affinity make sense physiologically?

The cognate vesicular transporter for GABA, VGAT, transports the inhibitory neurotransmitters GABA and glycine. Interestingly, the affinity of GABA for VGAT turned out to be surprisingly low (apparent K_M_ around 5 mM) (Burger et al., 1991; Hell et al., 1988; McIntire et al., 1997); and the affinity of glycine at VGAT was suggested to be even lower because it could not be measured experimentally (Edwards, 2007; McIntire et al., 1997). The affinity of GABA at VGAT is therefore considerably lower than the affinity of monoamines at VMAT2 (K_M_ values in the sub-micromolar range) (Liu et al., 1992; Yelin et al., 1995), but not so different from the affinity of glutamate at VGLUTs, which is also in the low millimolar range (Edwards, 2007; Maycox et al., 1988; Tabb et al., 1991).

Besides the affinity of a substrate to its transporter, the ambient concentration of the substrate determines whether it will be transported and concentrated within a compartment.

Cytosolic DA concentrations are estimated to be in the low micromolar range in adrenal chromaffin cells (Mosharov et al., 2003), and even lower in DA neurons (Mosharov et al., 2009) explaining why a sub-micromolar K_M_ of DA for VMAT2 allows for efficient sequestration and concentration of monoamines in synaptic vesicles at millimolar concentrations.

In contrast, cytosolic GABA concentrations remain unknown, but are presumably much higher than cytosolic monoamines (Edwards, 2007) and likely comparable to the estimated millimolar levels of glutamate (Featherstone, 2010; Ishikawa et al., 2002). Importantly, cytosolic GABA concentrations would need to be several-fold above the suggested K_M_ of 5 mM, to enable sufficient synaptic loading. If these concentrations are also reached within DA neurons (*e.g.* locally at uptake sites), GABA could be transported despite its apparent low affinity at VMAT2.

But is it really GABA that is transported by VMAT2 and released from DA neurons?

In fact, our evidence for GABA being a substrate at VMAT2 is indirect: we showed that increasing concentrations of GABA reduced the uptake of the VMAT2 substrate 5-HT into rodent synaptic vesicles. While we have attempted to directly measure uptake of [^3^H]GABA, we were not able to determine any reserpine- or tetrabenazine-sensitive VMAT2-mediated uptake of [^3^H]GABA into either rodent synaptic vesicles or VMAT2-expressing HEK-293 cells.

There are several possibilities that may explain these negative findings: *i)* GABA is not actually transported by VMAT2; *ii)* our uptake assays were not sensitive enough to measure uptake of a substrate with a low apparent affinity (similar to glycine at VGAT); *iii)* under physiological conditions, not GABA, but another inhibitory transmitter is taken up by VMAT2.

In fact, since GABA_A_ receptors are promiscuous ionotropic receptors that bind many different endogenous and exogenous ligands capable of activating or modulating the receptor, we tested two potential agonists that endogenously occur in mammals in high concentrations and have been shown to activate GABA_A_ receptors: glycine and taurine (Jonas et al., 1998). Interestingly, while glycine did not affect 5-HT uptake at VMAT2 in mouse synaptic vesicles, taurine weakly reduced 5-HT uptake, suggesting that it may also be taken up and released *in vivo*.

Our finding that GABA did not affect [^3^H]DA uptake in human VMAT2:eGFP-expressing HEK-293 cells could be explained by several factors: *i)* there could be species differences (mouse and human VMAT2 share 91 % of the amino acid sequence; mouse and rat VMAT2 share 95 %); *ii)* the C-terminal eGFP moiety may prevent the interaction of GABA and VMAT2; *iii)* the non-neuronal environment of HEK-293 cells lacks other important components (*e.g.* interacting proteins) only present in neuronal preparations.

Hence, we moved to synaptic vesicle preparations from rodent brains to determine whether GABA would reduce VMAT2 uptake and indeed found that GABA weakly reduced it. However, it is also important to stress that our synaptic vesicle preparation did not contain many of the additional features present in intact nerve cells including the surrounding boutons and local microdomains that may impact on transport. For instance, it is conceivable that *in vivo*, VMAT2 transport is further influenced by many additional factors such as the local concentrations of GABA, interacting proteins and differences in the local pH: the local concentration of GABA within specific boutons or presynaptic terminals could vary and therefore efficient uptake of GABA into vesicles may be enhanced, when local GABA concentrations are higher. VMAT2 and GAT1 may functionally couple as has been demonstrated for VGAT and GAD1/2 (Jin et al., 2003), or VMAT2 and the plasmalemmal dopamine transporter (DAT) (Cartier et al., 2010).

Finally, the local pH may play an important role in affecting the ionization of GABA (Krishek et al., 1996; Takeuchi et al., 1967). For instance, if the local pH is lower than the average cytosolic pH of 7.2, the relative stoichiometry of zwitterionic over cationic GABA may decrease, and affect uptake by VMAT2, which typically transports cations.

Lastly, our study demonstrates for the first time that not only mouse but also human midbrain DA neurons express GAT1 in about 50-60 % of the cells indicating that GABA co-release from DA neurons is evolutionarily conserved. In fact, the potential for GABA co-release from DA neurons may be far more prevalent than the co-release of glutamate which is restricted to only few DA neurons in SNc and VTA. Glutamate co-release from DA neurons has been implicated to play important roles in motivated behaviors and may also play a role in Parkinson’s disease (Hnasko et al., 2010; Shen et al., 2018; Steinkellner et al., 2022; Steinkellner et al., 2018). In contrast, the potential role of GABA co-release from DA neurons for physiology and disease is largely unknown though some evidence for a role in ethanol drinking and reward-based behaviors have been reported (Kim et al., 2015). Interestingly, we also found a subpopulation of DA neurons that was positive for VMAT2, GAT1 and VGLUT2, suggesting that these cells may be able to co-release three transmitters.

In conclusion, we provide pharmacological support that GABA could be a low-affinity atypical substrate at VMAT2, that seems to be taken up by GAT1 into the majority of SNc and many VTA DA neurons in mice and possibly also in humans.

## List of abbreviations

5HT: 5-hydroxytryptamine
DA: dopamine
DAT: dopamine transporter
DTBZ: dihydrotetrabenazine
GAD1/2: glutamic acid decarboxylase ½
GAT1: GABA transporter 1
NE: norepinephrine
SNc: substantia nigra pars compacta
TBZ: tetrabenazine
TH: tyrosine hydroxylase
VGAT: vesicular
GABA: transporter
VGLUT2: vesicular glutamate transporter 2
VMAT2: vesicular monoamine transporter 2
VTA: ventral tegmental area

## Data availability statement

The data that support the findings of this study are available from the corresponding author upon reasonable request. Some data may not be made available because of privacy or ethical restrictions.

## Declaration of transparency and scientific rigour

This declaration acknowledges that this paper adheres to the principles for transparent reporting and scientific rigour of preclinical research as stated in the BJP guidelines for Design and Analysis and as recommended by funding agencies, publishers, and other organisations engaged with supporting research.

## Ethics approval and consent to participate

All mice were used in accordance with protocols approved by the Animal Welfare

Committee of the Medical University of Vienna and the Austrian Federal Ministry of Science and Research (BMBWF licenses 2021-0.373.073 and 2023-0.515.074).

## Authors’ Contribution

FL, SS, MH, SLA and TS performed research. TS designed research with major input from TSH. TS acquired the funding. SK provided sections of human postmortem brains. TS wrote the manuscript with input from all authors. All authors read and approved the final manuscript.

## Acknowledgement

We thank Christian Pifl (Medical University of Vienna) for helpful tips regarding the vesicular uptake assays, Ellen Gelpi (Medical University of Vienna, Neurobiobank) for providing postmortem human samples and for initial neuropathological discussions and Stefan Boehm (Medical University of Vienna) for the female Sprague-Dawley rats.

We also thank the Core Facility Imaging of the Medical University of Vienna for their support. Most importantly, we are grateful to the patients who donated tissue to the neuro biobank at the Medical University of Vienna.

## Supplemental Figures

**Supplemental Figure 1:**
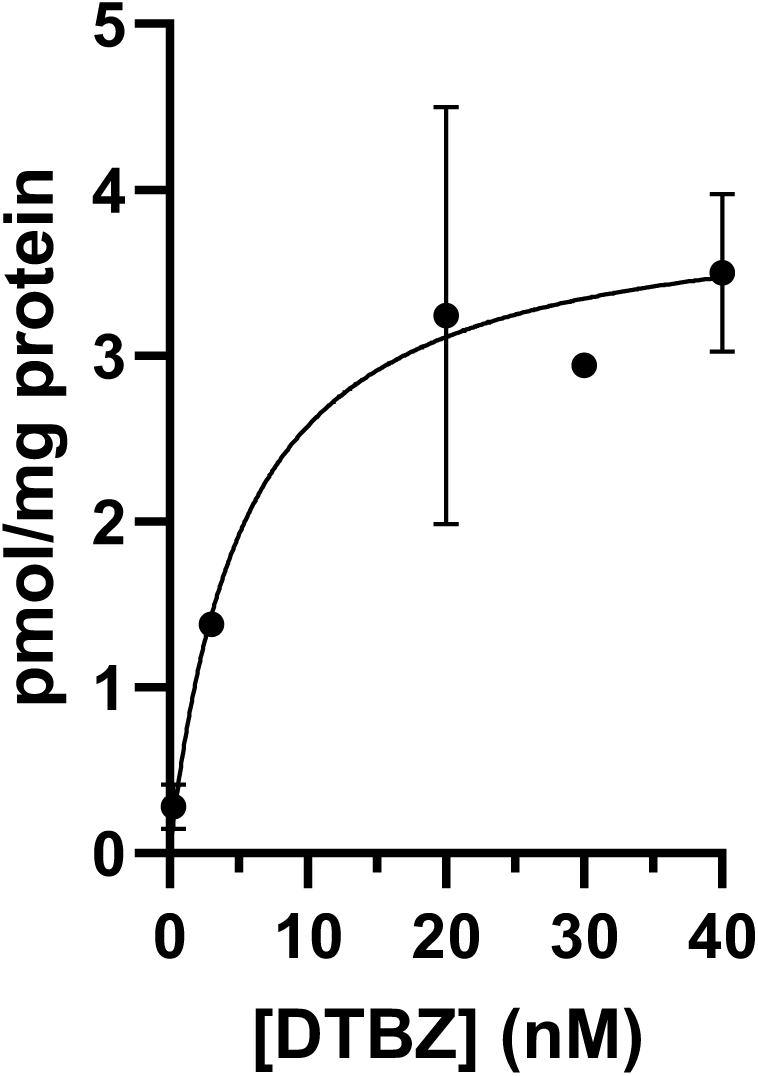
[^3^H]dihydrotetrabenazine binding in mouse striatal vesicles. Mouse striatal vesicles were incubated with increasing concentrations of [^3^H]dihydrotetrabenazine. Non-specific binding was performed in the presence of 10 µM unlabeled tetrabenazine. Experiments were performed in duplicate determinations from n=3 mice.

**Supplemental Figure 2:**
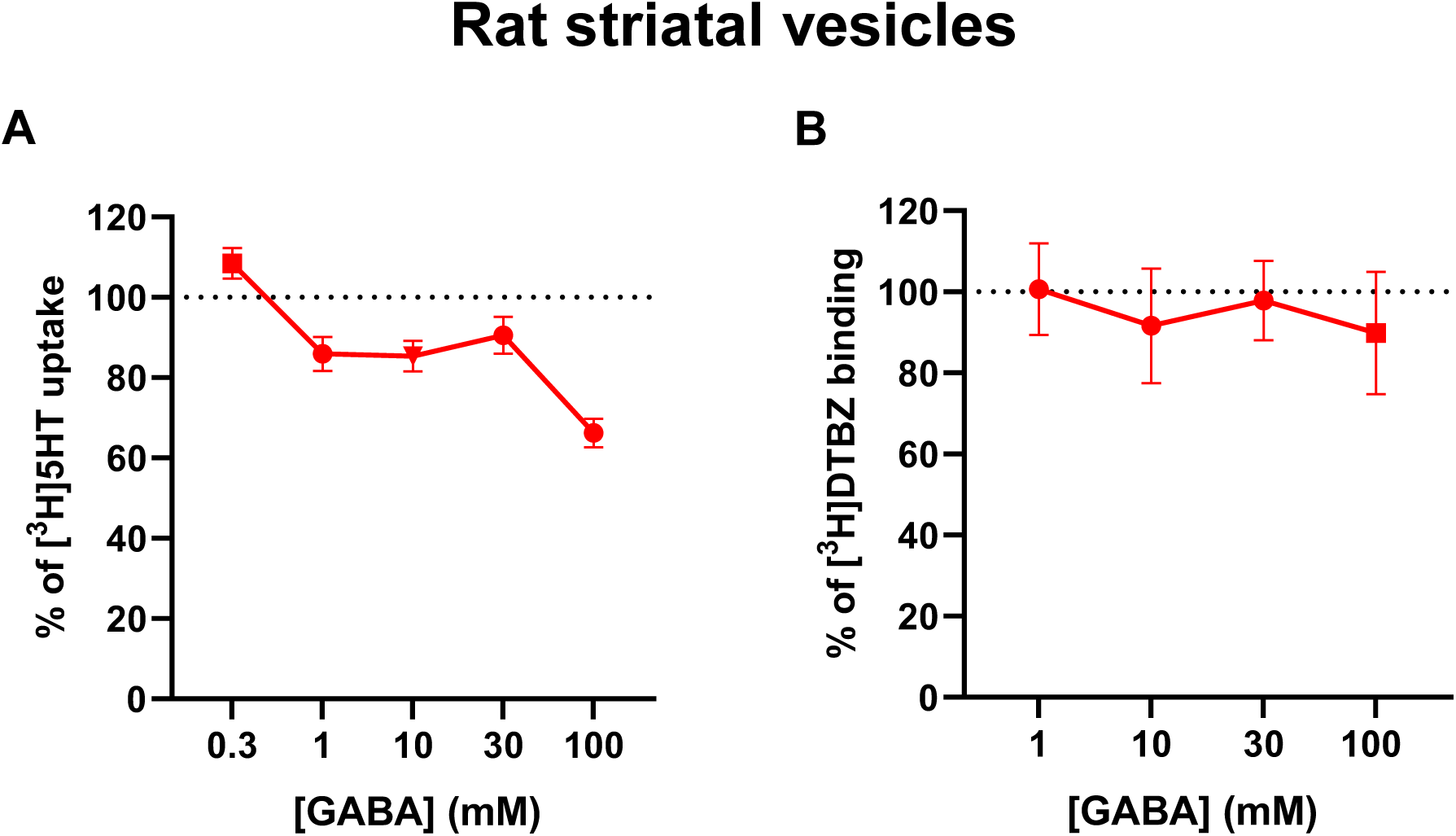
GABA decreased uptake of 5-HT in rat synaptic vesicles. **(A)** GABA concentration-dependently decreases [^3^H]5-HT uptake in striatal vesicles from rats. Uptakes were performed in duplicates prepared from the striata of n=6 female rats. **(B)** Binding of [^3^H]dihydrotetrabenazine is not affected by GABA in striatal synaptic vesicles; Experiments were performed in duplicate determinations from n=4 female rats.

**Supplemental Figure 3:**
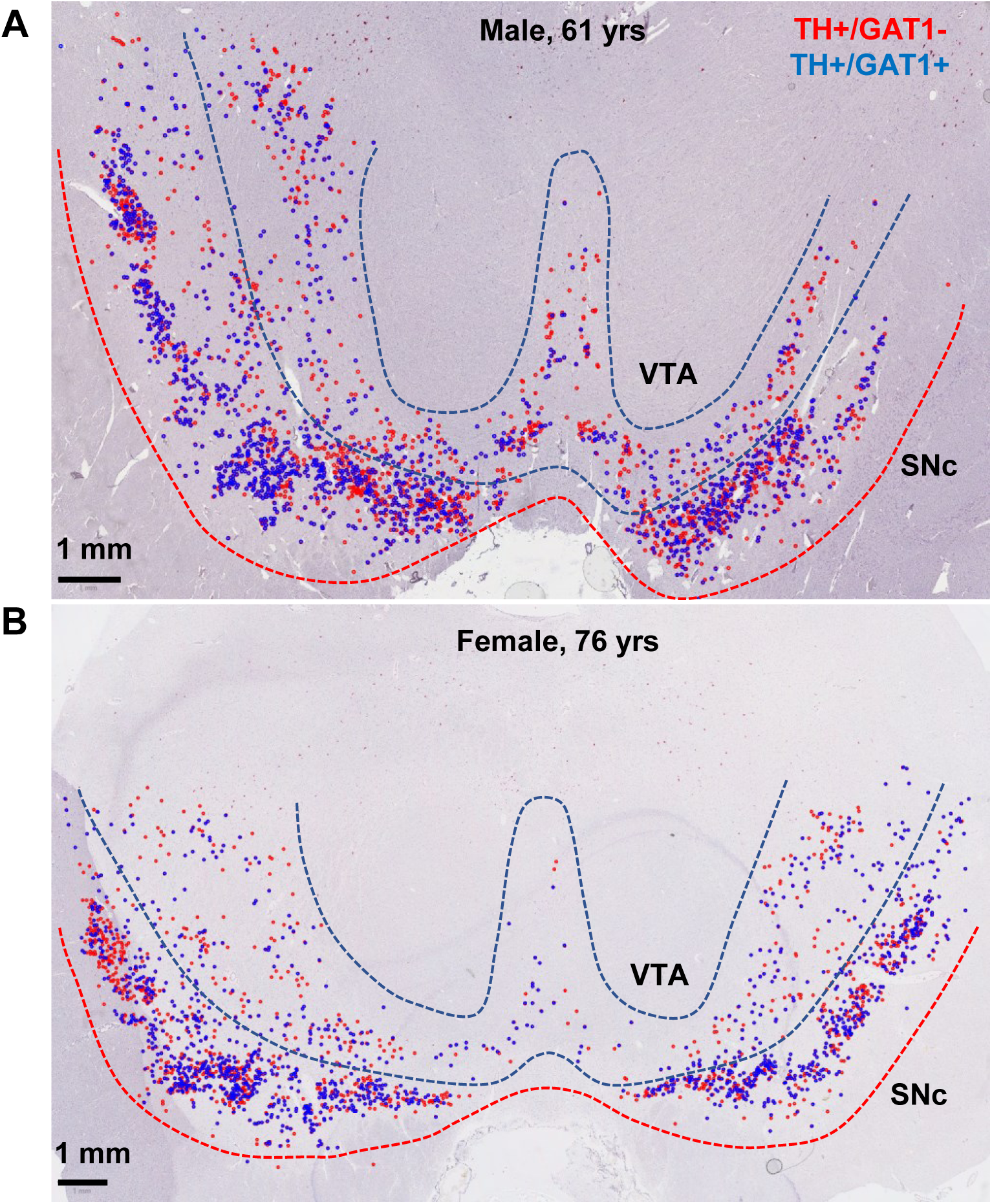
Overview of counting and distribution of counted cells in human midbrains. TH^+^/GAT1^+^ (blue dots) and TH^+^/GAT1^-^ (red dots) cells in the ventral midbrains of a male **(A)** and female **(B)** human donor.

## References

Adrover MF, Shin JH and Alvarez VA (2014) Glutamate and dopamine transmission from midbrain dopamine neurons share similar release properties but are differentially affected by cocaine. J Neurosci 34(9): 3183–3192. doi:10.1523/JNEUROSCI.4958-13.2014.

Azcorra M, Gaertner Z, et al. (2023) Unique functional responses differentially map onto genetic subtypes of dopamine neurons. Nat Neurosci 26(10): 1762–1774. doi:10.1038/s41593-023-01401-9.

Buck SA, Steinkellner T, et al. (2021) Vesicular glutamate transporter modulates sex differences in dopamine neuron vulnerability to age-related neurodegeneration. Aging Cell 20(5): e13365. doi:10.1111/acel.13365.

Burger PM, Hell J, et al. (1991) GABA and glycine in synaptic vesicles: storage and transport characteristics. Neuron 7(2): 287–293. doi:10.1016/0896-6273(91)90267-4.

Carlsson A (2001) A paradigm shift in brain research. Science 294(5544): 1021–1024. doi:10.1126/science.1066969.

Cartier EA, Parra LA, et al. (2010) A biochemical and functional protein complex involving dopamine synthesis and transport into synaptic vesicles. J Biol Chem 285(3): 1957–1966. doi:10.1074/jbc.M109.054510.

Chuhma N, Zhang H, et al. (2004) Dopamine neurons mediate a fast excitatory signal via their glutamatergic synapses. J Neurosci 24(4): 972–981. doi:10.1523/JNEUROSCI.4317-03.2004.

Conrad WS, Oriol L, Faget L and Hnasko TS (2024) Proportion and distribution of neurotransmitter-defined cell types in the ventral tegmental area and substantia nigra pars compacta. bioRxiv. doi:10.1101/2024.02.28.582356.

Costa KM and Schoenbaum G (2022) Dopamine. Curr Biol 32(15): R817–R824. doi:10.1016/j.cub.2022.06.060.

Dahlstroem A and Fuxe K (1964) Evidence for the Existence of Monoamine-Containing Neurons in the Central Nervous System. I. Demonstration of Monoamines in the Cell Bodies of Brain Stem Neurons. Acta Physiol Scand Suppl: SUPPL 232:231–255.

Dal Bo G, Berube-Carriere N, et al. (2008) Enhanced glutamatergic phenotype of mesencephalic dopamine neurons after neonatal 6-hydroxydopamine lesion. Neuroscience 156(1): 59–70. doi:10.1016/j.neuroscience.2008.07.032.

Damier P, Hirsch EC, Agid Y and Graybiel AM (1999) The substantia nigra of the human brain. II. Patterns of loss of dopamine-containing neurons in Parkinson’s disease. Brain 122 ( **Pt 8**): 1437–1448. doi:10.1093/brain/122.8.1437.

Edwards RH (2007) The neurotransmitter cycle and quantal size. Neuron 55(6): 835–858. doi:10.1016/j.neuron.2007.09.001.

Eiden LE and Weihe E (2011) VMAT2: a dynamic regulator of brain monoaminergic neuronal function interacting with drugs of abuse. Ann N Y Acad Sci 1216: 86–98. doi:10.1111/j.1749-6632.2010.05906.x.

Erickson JD, Eiden LE and Hoffman BJ (1992) Expression cloning of a reserpine-sensitive vesicular monoamine transporter. Proc Natl Acad Sci U S A 89(22): 10993–10997. doi:10.1073/pnas.89.22.10993.

Featherstone DE (2010) Intercellular glutamate signaling in the nervous system and beyond. ACS Chem Neurosci 1(1): 4–12. doi:10.1021/cn900006n.

Gaertner Z, Oram C, et al. (2024) Molecular and spatial transcriptomic classification of midbrain dopamine neurons and their alterations in a LRRK2(G2019S) model of Parkinson’s disease. bioRxiv. doi:10.1101/2024.06.06.597807.

German DC, Nelson EL, et al. (1996) The neurotoxin MPTP causes degeneration of specific nucleus A8, A9 and A10 dopaminergic neurons in the mouse. Neurodegeneration 5(4): 299–312. doi:10.1006/neur.1996.0041.

Glaser PE, Surgener SP, et al. (2006) Cerebellar neurotransmission in attention-deficit/hyperactivity disorder: does dopamine neurotransmission occur in the cerebellar vermis? J Neurosci Methods 151(1): 62–67. doi:10.1016/j.jneumeth.2005.09.019.

Hell JW, Maycox PR, Stadler H and Jahn R (1988) Uptake of GABA by rat brain synaptic vesicles isolated by a new procedure. EMBO J 7(10): 3023–3029. doi:10.1002/j.1460-2075.1988.tb03166.x.

Hirsch EC, Graybiel AM and Agid Y (1989) Selective vulnerability of pigmented dopaminergic neurons in Parkinson’s disease. Acta Neurol Scand Suppl 126: 19–22. doi:10.1111/j.1600-0404.1989.tb01778.x.

Hnasko TS, Chuhma N, et al. (2010) Vesicular glutamate transport promotes dopamine storage and glutamate corelease in vivo. Neuron 65(5): 643–656. doi:10.1016/j.neuron.2010.02.012.

Hornykiewicz O (2006) The discovery of dopamine deficiency in the parkinsonian brain. J Neural Transm Suppl(70): 9–15. doi:10.1007/978-3-211-45295-0_3.

Ishikawa T, Sahara Y and Takahashi T (2002) A single packet of transmitter does not saturate postsynaptic glutamate receptors. Neuron 34(4): 613–621. doi:10.1016/s0896-6273(02)00692-x.

Iversen SD and Iversen LL (2007) Dopamine: 50 years in perspective. Trends Neurosci 30(5): 188–193. doi:10.1016/j.tins.2007.03.002.

Jin H, Wu H, et al. (2003) Demonstration of functional coupling between gamma-aminobutyric acid (GABA) synthesis and vesicular GABA transport into synaptic vesicles. Proc Natl Acad Sci U S A 100(7): 4293–4298. doi:10.1073/pnas.0730698100.

Jonas P, Bischofberger J and Sandkuhler J (1998) Corelease of two fast neurotransmitters at a central synapse. Science 281(5375): 419–424. doi:10.1126/science.281.5375.419.

Kanner BI, Fishkes H, Maron R, Sharon I and Schuldiner S (1979) Reserpine as a competitive and reversible inhibitor of the catecholamine transporter of bovine chromaffin granules. FEBS Lett 100(1): 175–178. doi:10.1016/0014-5793(79)81158-8.

Kawano M, Kawasaki A, et al. (2006) Particular subpopulations of midbrain and hypothalamic dopamine neurons express vesicular glutamate transporter 2 in the rat brain. J Comp Neurol 498(5): 581–592. doi:10.1002/cne.21054.

Kim JI, Ganesan S, et al. (2015) Aldehyde dehydrogenase 1a1 mediates a GABA synthesis pathway in midbrain dopaminergic neurons. Science 350(6256): 102–106. doi:10.1126/science.aac4690.

Krishek BJ, Amato A, Connolly CN, Moss SJ and Smart TG (1996) Proton sensitivity of the GABA(A) receptor is associated with the receptor subunit composition. J Physiol 492 (**Pt 2**)(Pt 2): 431–443. doi:10.1113/jphysiol.1996.sp021319.

Liu Y, Peter D, et al. (1992) A cDNA that suppresses MPP+ toxicity encodes a vesicular amine transporter. Cell 70(4): 539–551. doi:10.1016/0092-8674(92)90425-c.

Maycox PR, Deckwerth T, Hell JW and Jahn R (1988) Glutamate uptake by brain synaptic vesicles. Energy dependence of transport and functional reconstitution in proteoliposomes. J Biol Chem 263(30): 15423–15428.

McIntire SL, Reimer RJ, Schuske K, Edwards RH and Jorgensen EM (1997) Identification and characterization of the vesicular GABA transporter. Nature 389(6653): 870–876. doi:10.1038/39908.

Melani R and Tritsch NX (2022) Inhibitory co-transmission from midbrain dopamine neurons relies on presynaptic GABA uptake. Cell Rep 39(3): 110716. doi:10.1016/j.celrep.2022.110716.

Mosharov EV, Gong LW, Khanna B, Sulzer D and Lindau M (2003) Intracellular patch electrochemistry: regulation of cytosolic catecholamines in chromaffin cells. J Neurosci 23(13): 5835–5845. doi:10.1523/JNEUROSCI.23-13-05835.2003.

Mosharov EV, Larsen KE, et al. (2009) Interplay between cytosolic dopamine, calcium, and alpha-synuclein causes selective death of substantia nigra neurons. Neuron 62(2): 218–229. doi:10.1016/j.neuron.2009.01.033.

Nelson EL, Liang CL, Sinton CM and German DC (1996) Midbrain dopaminergic neurons in the mouse: computer-assisted mapping. J Comp Neurol 369(3): 361–371. doi:10.1002/(SICI)1096-9861(19960603)369:3<361::AID-CNE3>3.0.CO;2-3.

Patel JC, Sherpa AD, et al. (2024) GABA co-released from striatal dopamine axons dampens phasic dopamine release through autoregulatory GABA(A) receptors. Cell Rep 43(3): 113834. doi:10.1016/j.celrep.2024.113834.

Peneder TM, Scholze P, et al. (2011) Chronic exposure to manganese decreases striatal dopamine turnover in human alpha-synuclein transgenic mice. Neuroscience 180: 280–292. doi:10.1016/j.neuroscience.2011.02.017.

Peter D, Jimenez J, Liu Y, Kim J and Edwards RH (1994) The chromaffin granule and synaptic vesicle amine transporters differ in substrate recognition and sensitivity to inhibitors. J Biol Chem 269(10): 7231–7237.

Pifl C, Rajput A, et al. (2014) Is Parkinson’s disease a vesicular dopamine storage disorder? Evidence from a study in isolated synaptic vesicles of human and nonhuman primate striatum. J Neurosci 34(24): 8210–8218. doi:10.1523/JNEUROSCI.5456-13.2014.

Poulin JF, Gaertner Z, Moreno-Ramos OA and Awatramani R (2020) Classification of Midbrain Dopamine Neurons Using Single-Cell Gene Expression Profiling Approaches. Trends Neurosci 43(3): 155–169. doi:10.1016/j.tins.2020.01.004.

Root DH, Wang HL, et al. (2016) Glutamate neurons are intermixed with midbrain dopamine neurons in nonhuman primates and humans. Sci Rep 6: 30615. doi:10.1038/srep30615.

Shen H, Marino RAM, et al. (2018) Genetic deletion of vesicular glutamate transporter in dopamine neurons increases vulnerability to MPTP-induced neurotoxicity in mice. Proc Natl Acad Sci U S A 115(49): E11532–E11541. doi:10.1073/pnas.1800886115.

Silm K, Yang J, et al. (2019) Synaptic Vesicle Recycling Pathway Determines Neurotransmitter Content and Release Properties. Neuron 102(4): 786–800 e785. doi:10.1016/j.neuron.2019.03.031.

Steinkellner T, Conrad WS, et al. (2022) Dopamine neurons exhibit emergent glutamatergic identity in Parkinson’s disease. Brain 145(3): 879–886. doi:10.1093/brain/awab373.

Steinkellner T, Zell V, et al. (2018) Role for VGLUT2 in selective vulnerability of midbrain dopamine neurons. J Clin Invest 128(2): 774–788. doi:10.1172/JCI95795.

Stuber GD, Hnasko TS, Britt JP, Edwards RH and Bonci A (2010) Dopaminergic terminals in the nucleus accumbens but not the dorsal striatum corelease glutamate. J Neurosci 30(24): 8229–8233. doi:10.1523/JNEUROSCI.1754-10.2010.

Tabb JS and Ueda T (1991) Phylogenetic studies on the synaptic vesicle glutamate transport system. J Neurosci 11(6): 1822–1828. doi:10.1523/JNEUROSCI.11-06-01822.1991.

Takamori S, Rhee JS, Rosenmund C and Jahn R (2000) Identification of a vesicular glutamate transporter that defines a glutamatergic phenotype in neurons. Nature 407(6801): 189–194. doi:10.1038/35025070.

Takeuchi A and Takeuchi N (1967) Anion permeability of the inhibitory post-synaptic membrane of the crayfish neuromuscular junction. J Physiol 191(3): 575–590. doi:10.1113/jphysiol.1967.sp008269.

Tritsch NX, Ding JB and Sabatini BL (2012) Dopaminergic neurons inhibit striatal output through non-canonical release of GABA. Nature 490(7419): 262–266. doi:10.1038/nature11466.

Tritsch NX, Granger AJ and Sabatini BL (2016) Mechanisms and functions of GABA co-release. Nat Rev Neurosci 17(3): 139–145. doi:10.1038/nrn.2015.21.

Tritsch NX, Oh WJ, Gu C and Sabatini BL (2014) Midbrain dopamine neurons sustain inhibitory transmission using plasma membrane uptake of GABA, not synthesis. Elife 3: e01936. doi:10.7554/eLife.01936.

Trudeau LE, Hnasko TS, et al. (2014) The multilingual nature of dopamine neurons. Prog Brain Res 211: 141–164. doi:10.1016/B978-0-444-63425-2.00006-4.

Wallace ML and Sabatini BL (2023) Synaptic and circuit functions of multitransmitter neurons in the mammalian brain. Neuron 111(19): 2969–2983. doi:10.1016/j.neuron.2023.06.003.

Yamaguchi T, Wang HL, Li X, Ng TH and Morales M (2011) Mesocorticolimbic glutamatergic pathway. J Neurosci 31(23): 8476–8490. doi:10.1523/JNEUROSCI.1598-11.2011.

Yelin R and Schuldiner S (1995) The pharmacological profile of the vesicular monoamine transporter resembles that of multidrug transporters. FEBS Lett 377(2): 201–207. doi:10.1016/0014-5793(95)01346-6.

Zheng G, Dwoskin LP and Crooks PA (2006) Vesicular monoamine transporter 2: role as a novel target for drug development. AAPS J 8(4): E682–692. doi:10.1208/aapsj080478.

